# Evolution of a novel chimeric maltotriose transporter in *Saccharomyces eubayanus* from parent proteins unable to perform this function

**DOI:** 10.1101/431171

**Authors:** EmilyClare P. Baker, Chris Todd Hittinger

## Abstract

At the molecular level, the evolution of new traits can be broadly divided between changes in gene expression and changes in protein structure. For proteins, the evolution of novel functions is generally thought to proceed through sequential point mutations or recombination of whole functional units. In *Saccharomyces*, the uptake of the sugar maltotriose into the cell is the primary limiting factor in its utilization, but maltotriose transporters are relatively rare, except in brewing strains. No known wild strains of *Saccharomyces eubayanus*, the cold-tolerant parent of hybrid lager-brewing yeasts (*Saccharomyces cerevisiae x S. eubayanus*), are able to consume maltotriose, which limits their ability to fully ferment malt extract. In one strain of *S. eubayanus*, we found a gene closely related to a known maltotriose transporter and were able to confer maltotriose consumption by overexpressing this gene or by passaging the strain on maltose. Even so, most wild strains of *S*. *eubayanus* lack native maltotriose transporters. To determine how this rare trait could evolve in naive genetic backgrounds, we performed an adaptive evolution experiment for maltotriose consumption, which yielded a single strain of *S*. *eubayanus* able to grow on maltotriose. We mapped the causative locus to a gene encoding a novel chimeric transporter that was formed by an ectopic recombination event between two genes encoding transporters that are unable to import maltotriose. In contrast to classic models of the evolution of novel protein functions, the recombination breakpoints occurred within functional domains. Thus, the ability of the new protein to carry maltotriose was likely acquired through epistatic interactions between independently evolved substitutions. By acquiring multiple mutations at once, the transporter rapidly gained a novel function, while bypassing potentially deleterious intermediate steps. This study provides an illuminating example of how recombination between paralogs can establish novel interactions among substitutions to create adaptive functions.

**Author summary:** Hybrids of the yeasts *Saccharomyces cerevisiae* and *Saccharomyces eubayanus* (lager-brewing yeasts) dominate the modern brewing industry. *S*. *cerevisiae*, also known as baker’s yeast, is well-known for its role in industry and scientific research. Less well recognized is *S*. *eubayanus*, which was only discovered as a pure species in 2011. While most lager-brewing yeasts rapidly and completely utilize the important brewing sugar maltotriose, no strain of *S*. *eubayanus* isolated to date is known to do so. Despite being unable to consume maltotriose, we identified one strain of *S*. *eubayanus* carrying a gene for a functional maltotriose transporter, although most strains lack this gene. During an adaptive evolution experiment, a strain of *S*. *eubayanus* without native maltotriose transporters evolved the ability to grow on maltotriose. Maltotriose consumption in the evolved strain resulted from a chimeric transporter that arose through recombination between genes encoding parent proteins that were unable to transport maltotriose. Traditionally, functional chimeric proteins are thought to evolve by recombining discrete functional domains or modules, but the breakpoints in the chimera studied here occurred within modular units of the protein. These results support the less well-recognized role of recombination between paralogous sequences in generating novel proteins with adaptive functions.

## Introduction

Proteins with novel functions can arise through a variety of mechanisms (1). One of the best studied mechanisms is gene duplication, followed by divergence through sequential point mutations (1,2). While this method of new protein evolution is thought to be common, evolution through stepwise point mutations can be a slow and constrained process (3). In the mutational landscape separating the original protein from the derived protein, deleterious epistatic interactions, where multiple intermediate mutational steps interact to create fitness valleys, can make new functions difficult to access by successive point mutations. Mutational events that result in multiple amino acid changes at once can help bridge fitness valleys and speed the evolution of new functionality (3,4). As a consequence of bypassing intermediate mutational steps, recombination can lead to intragenic reciprocal sign epistasis, where the new recombinant protein has a function not found in either parent protein (3,5). Ectopic gene conversion, which results in chimeric protein sequences, is one such rare class of mutational events that can rapidly lead to new protein sequences with novel functions (3).

Chimeric protein-coding sequences have been found to be an important mechanism by which proteins can evolve new functions (1,6–8). They have been implicated in the rapid radiation of multicellular animals (6) and in playing a role in both infectious and non-infectious diseases in humans (9–12). The *Drosophila* gene *jingwei* was one of the first chimeric genes to have both its recent origin and evolution characterized in depth (1). *jingwei* exemplifies many of the characteristics usually associated with chimeric proteins (1,6–8,13). Like most other chimeric proteins that have been described in eukaryotes, *jingwei* is a large multidomain protein that was constructed via the movement of whole functional units (domains), facilitated by intronic sequences, a process referred to as domain or exon shuffling. In most cases, even in the absence of intronic sequences, the recombination of these modules has been considered key to the evolution of functional chimeric proteins (1,14).

The exchange of complete, independently functional units is not the only method by which functional chimeric proteins can be generated. Recombination within functional domains also has the potential to create proteins with novel characteristics. Recombination breakpoints within domains can lead to functional proteins, even between non-homologous protein sequences (4,15,16). However, since functionally important structures are likely to be conserved between related proteins, the probability of recombination resulting in a functional protein is higher between homologous sequences where essential within-protein interactions are less likely to be disrupted (4,15,17). Theoretical work has suggested the potential of this sort of recombination to allow proteins to rapidly bypass fitness minima in the adaptive landscape separating two protein functions (3,4). Recombination between paralogous sequences has also been shown to be selected for in natural populations, suggesting that such sequences can indeed produce functional proteins (15,18). In addition, recombination between paralogous sequences (DNA shuffling) has been used as an efficient way to engineer proteins with functions that are rare or difficult to evolve in natural settings (reviewed in (19–21)). For example, hexose sugar transporters in *Saccharomyces cerevisiae* were evolved for increased specificity to a pentose sugar, D-xylose (22), through DNA shuffling and selection for the ability to support growth on xylose.

Maltotriose, a trimer of glucose molecules, is the second most abundant fermentable sugar present in brewing wort (malt extract), but it is also the most difficult to ferment (23–26). Among budding yeasts of the genus *Saccharomyces*, such as *S*. *cerevisiae*, proteins that can transport maltotriose into the cell are relatively rare (27–31). Improving consumption of maltotriose by *Saccharomyces* yeasts is of general interest to the brewing community since a key consideration for any new brewing strain is its ability to rapidly and completely ferment all the sugars present in wort. Work on improving the direct uptake of maltotriose in brewing yeasts has focused on the expression of the limited set of known maltotriose transporters, either through adaptive evolution for increased expression (32,33), introducing maltotriose transporters into new strains through selective breeding (34–42), or by heterologous expression (33,40,43). These methods all rely on the presence of functional maltotriose transporters, either natively or heterologously expressed, and are limited by the number of strains and proteins that are known to be capable of transporting maltotriose. With the focus on known transporters, how new maltotriose transporters evolve is less well studied (44,45).

Recently, special interest has been given to the development of *Saccharomyces eubayanus*, a distant cold-tolerant relative of *S*. *cerevisiae*, for commercial brewing (35,36,46,47). As a hybrid with *S*. *cerevisiae, S. eubayanus* forms the industrially important lager-brewing yeasts (48), which account for more than 90% of the total beer market. So far, no strain of *S*. *eubayanus* isolated from nature has been reported to consume maltotriose (36,49–53), despite evidence for the possible presence of functional transporters in the *S. eubayanus* subgenome of industrial *S. cerevisiae* x *S. eubayanus* hybrids (i.e. lager-brewing yeasts) (28,54–58).

In the present study, we characterize the native *MALT* genes found in *S. eubayanus* for their ability to enable the transport of maltotriose and confirm the presence of maltotriose transporters in one strain of *S. eubayanus*, despite its inability to consume maltotriose. We also describe a novel chimeric maltotriose transporter that resulted from the adaptive evolution of *S. eubayanus* for maltotriose consumption. This new maltotriose transporter was formed through a partial ectopic gene conversion event between two *MALT* genes. Interestingly, the parent proteins that produced the chimera were unable to transport maltotriose themselves. In addition, the breakpoints of the chimeric region do not demarcate clearly defined functional domains, suggesting that epistatic interactions between novel residue combinations, rather than domain swapping, is responsible for the new function. Overall, this study reports the first known maltotriose transporters in *S. eubayanus* and the first strains of this species that are able to consume maltotriose. In addition, by characterizing one of the few chimeric proteins that have been described where recombination occurred naturally within functional modules (without being specifically targeted for engineering by DNA shuffling or mutagenesis), we provide insight into how proteins can evolve novel adaptive functions through rare genetic events.

## Results/Discussion

### Maltotriose transporters in *S. eubayanus*

In the type strain of *S*. eubayanus, four genes, designated *MALT1-4*, have been identified as having homology to genes encoding known maltose transporters (*MALT* genes) (54,59). Because *MALT2* and *MALT4* are predicted to encode identical amino acid sequences (see Materials and Methods), we refer to these genes jointly as *MALT2/4*. To determine if they could enable maltotriose transport, Malt1, Malt2/4, and Malt3 were individually overexpressed using an inducible promoter in yHRVM108, a strain of *S. eubayanus* isolated from North Carolina that is unable to grow on maltotriose and, unlike other strains of *S. eubayanus*, grows sluggishly on maltose. None of these genes were able to confer growth on maltotriose when overexpressed (Table 1).

**Table 1.**
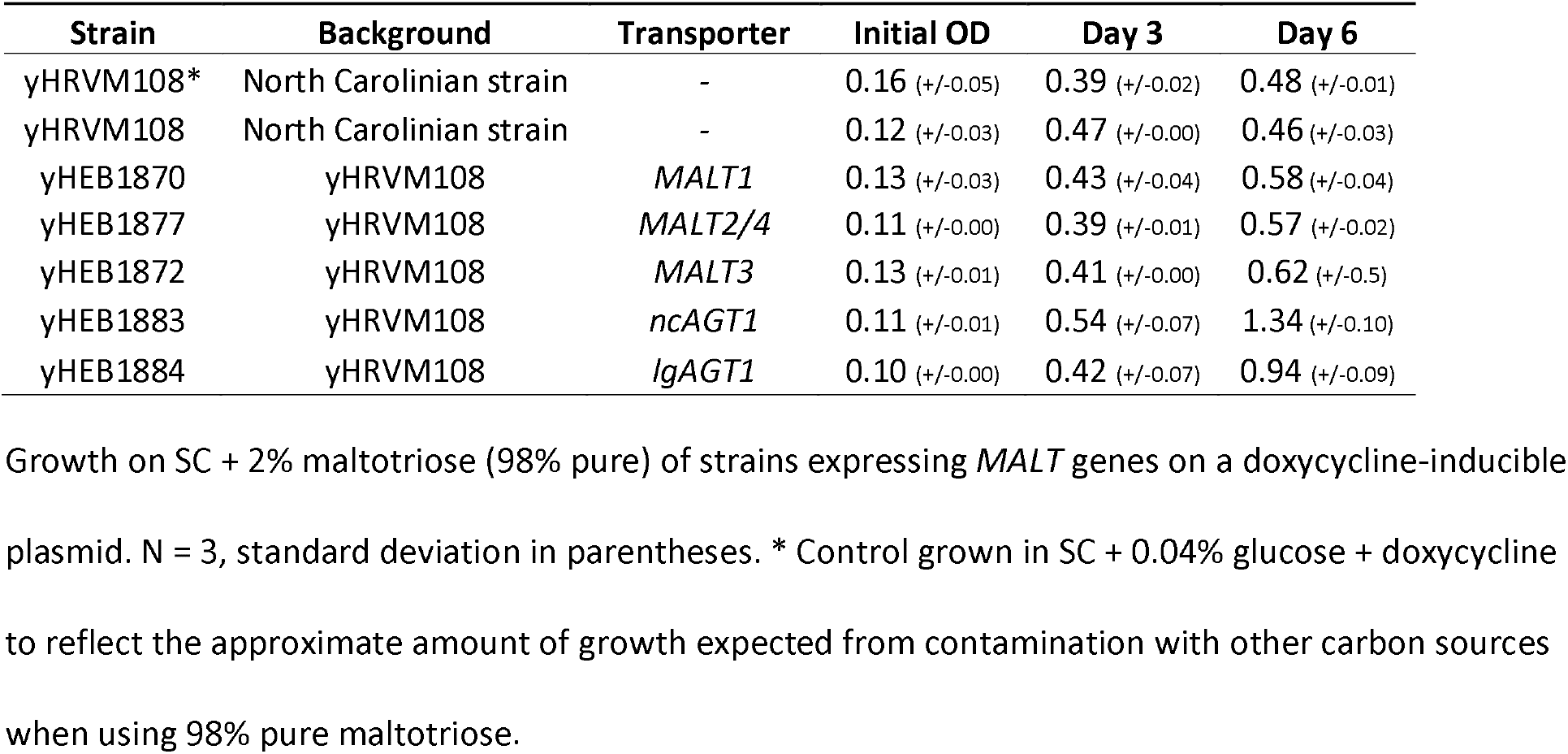
Heterologous expression of *S. eubayanus MALT* genes.

| Strain | Background | Transporter | Initial OD | Day 3 | Day 6 |
| --- | --- | --- | --- | --- | --- |
| yHRVM108* | North Carolinian strain | - | 0.16 (+/-0.05) | 0.39 (+/-0.02) | 0.48 (+/-0.01) |
| yHRVM108 | North Carolinian strain | - | 0.12 (+/-0.03) | 0.47 (+/-0.00) | 0.46 (+/-0.03) |
| yHEB1870 | yHRVM108 | *MALT1* | 0.13 (+/-0.03) | 0.43 (+/-0.04) | 0.58 (+/-0.04) |
| yHEB1877 | yHRVM108 | *MALT2/4* | 0.11 (+/-0.00) | 0.39 (+/-0.01) | 0.57 (+/-0.02) |
| yHEB1872 | yHRVM108 | *MALT3* | 0.13 (+/-0.01) | 0.41 (+/-0.00) | 0.62 (+/-0.5) |
| yHEB1883 | yHRVM108 | *ncAGT1* | 0.11 (+/-0.01) | 0.54 (+/-0.07) | 1.34 (+/-0.10) |
| yHEB1884 | yHRVM108 | *lgAGT1* | 0.10 (+/-0.00) | 0.42 (+/-0.07) | 0.94 (+/-0.09) |

Growth on SC + 2% maltotriose (98% pure) of strains expressing *MALT* genes on a doxycycline-inducible plasmid. N = 3, standard deviation in parentheses. \* Control grown in SC + 0.04% glucose + doxycycline to reflect the approximate amount of growth expected from contamination with other carbon sources when using 98% pure maltotriose.

Although none of the transporters found in the type strain of *S. eubayanus* were able to support growth on maltotriose, there is compelling evidence from lager-brewing yeasts for the existence of maltotriose transporters within the greater *S*. *eubayanus* population (28,54–57). Of particular interest are alleles of *AGT1*. Two versions of *AGT1* are present in the genomes of lager-brewing yeasts. One, which we call *scAGT1* (*S. cerevisiae-AGT1*), was donated by the *S. cerevisiae* parent of lager yeasts, and the other, which we call *lgAGT1* (lager*-AGT1*), has been proposed to be of *S. eubayanus* origin (55). Both *lgAGT1* and *scAGT1*, like other *AGT1* alleles, can transport both maltose and maltotriose (27,28,57,60–63). Thus far, full-length sequences closely related to this *lgAGT1* have not been described in any strain of *S. eubayanus* (36).

Strain CDFM21L.1 and a closely related strain isolated from North Carolina, yHRVM108, belong to the Holarctic subpopulation of *S. eubayanus* and are close relatives of the strains of *S. eubayanus* that hybridized with *S. cerevisiae* to form lager-brewing yeasts (52). Because of their close phylogenetic relationship, CDFM21L.1, yHRVM108, and the *S. eubayanus* lager parent are more likely to share strain-specific genes, such as *lgAGT1* (64). From a search of Illumina sequencing reads available for CDFM21L.1 and yHRVM108, we were able to assemble two full-length genes with high sequence identity to *lgAGT1*, which we designated *tbAGT1* and *ncAGT1*, for Tibetan-*AGT1* and North Carolinian-*AGT1*, respectively (Fig 1).

**Fig 1.**
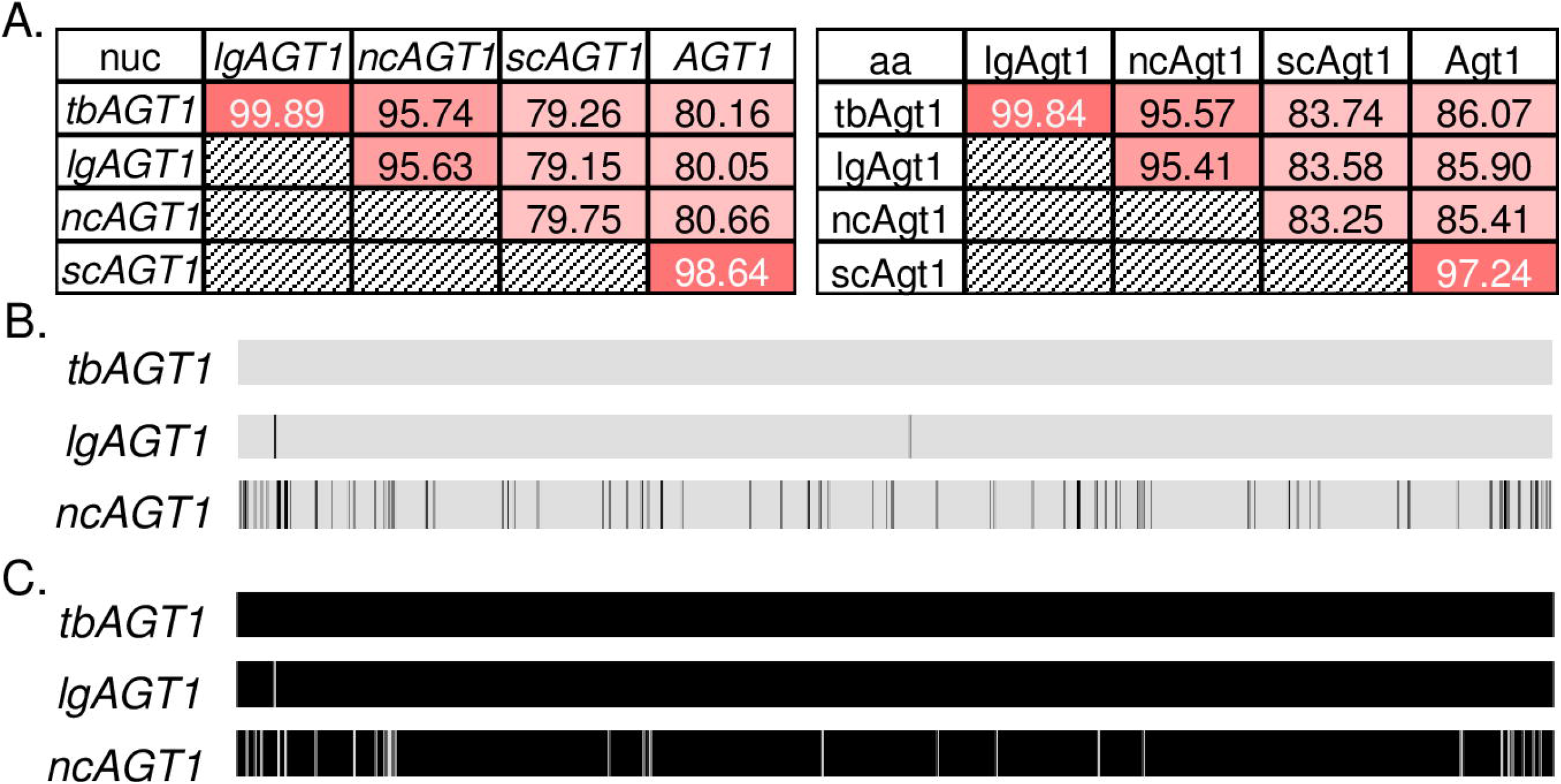
Alignment of *AGT1*-like genes. A) Tables highlighting the nucleotide (nuc) and amino acid (aa) percent identities between members of the *AGT1* family. Darker colors indicate greater sequence similarity. B) Multiple sequence alignment between nucleotide sequences of *tbAGT1, lgAGT1*, and *ncAGT1*. Black lines indicate nucleotide differences. C) Multiple sequence alignments between protein sequences of *tbAGT1, lgAGT1*, and *ncAGT1*. White gaps indicate amino acid differences.

Two single nucleotide polymorphisms (SNPs) separate *tbAGT1* and *lgAGT1*. One SNP results in a synonymous substitution and the other in a nonsynonymous substitution near the N-terminus of the protein outside of any predicted transmembrane domains (Fig 1B and C, S1 Fig). Analyses of the predicted effect of this substitution in *lgAGT1* (using STRUM and SIFT mutant protein prediction software (65,66)) suggest that it is unlikely to significantly impact protein structure or function (S1 Table). In contrast, *ncAGT1* has 95% nucleotide identity with *lgAGT1*, with nonsynonymous differences distributed throughout the sequence (Fig 1A-C). Despite the presence of *ncAGT1*, the yHRVM108 wild-type strain grows poorly on maltose and is unable to grow on maltotriose. Interestingly, and unlike all *MALT* genes found in the Patagonian type strain of *S. eubayanus*, overexpression of *ncAGT1* in yHRVM108 conferred growth on maltotriose, similar to the known maltotriose transporter gene *lgAGT1* (Table 1). These results suggest that insufficient *ncAGT1* gene expression, rather than protein function, is likely the main reason for the inability of yHRVM108 to grow on maltotriose.

### Phylogenetic relationships among maltose and maltotriose transporters

To put the relationship between *S. eubayanus, S. cerevisiae*, and lager *MALT* genes into a phylogenetic perspective, a gene tree was constructed for these three groups of genes (Fig 2). Consistent with previous analyses of *MALT* genes in *Saccharomyces* (30), the *MALT* genes fell into 3 major clades. The *AGT1* genes formed their own group, significantly divergent from the other clades and was further split between the *AGT1* genes originating from *S. cerevisiae* and the *AGT1* genes originating from *S. eubayanus. MPH* genes, which are native to *S. cerevisiae* but also present in some lager yeasts (63,67), also formed their own clade. *MPH* genes are most often described as encoding maltose transporters, but their ability to transport maltotriose is still uncertain (30,63,67–69). The final and largest clade was made up of *MALT1-4* from *S. eubayanus, MALx1* genes from *S. cerevisiae*, and the lager-specific gene *MTT1* (28,29,54). This clade was further subdivided into a group containing only *S. eubayanus MALT* genes and their close lager homologs and another group consisting of *MALx1* genes, *MTT1*, and *MALT3*. Within this clade, genes encoding maltotriose transporters were rare, represented by only a single gene, *MTT1* (28,29). The phylogenetic distribution of maltotriose utilization suggests that the ability to transport maltotriose may be a difficult function for genes within this clade to evolve.

**Fig 2.**
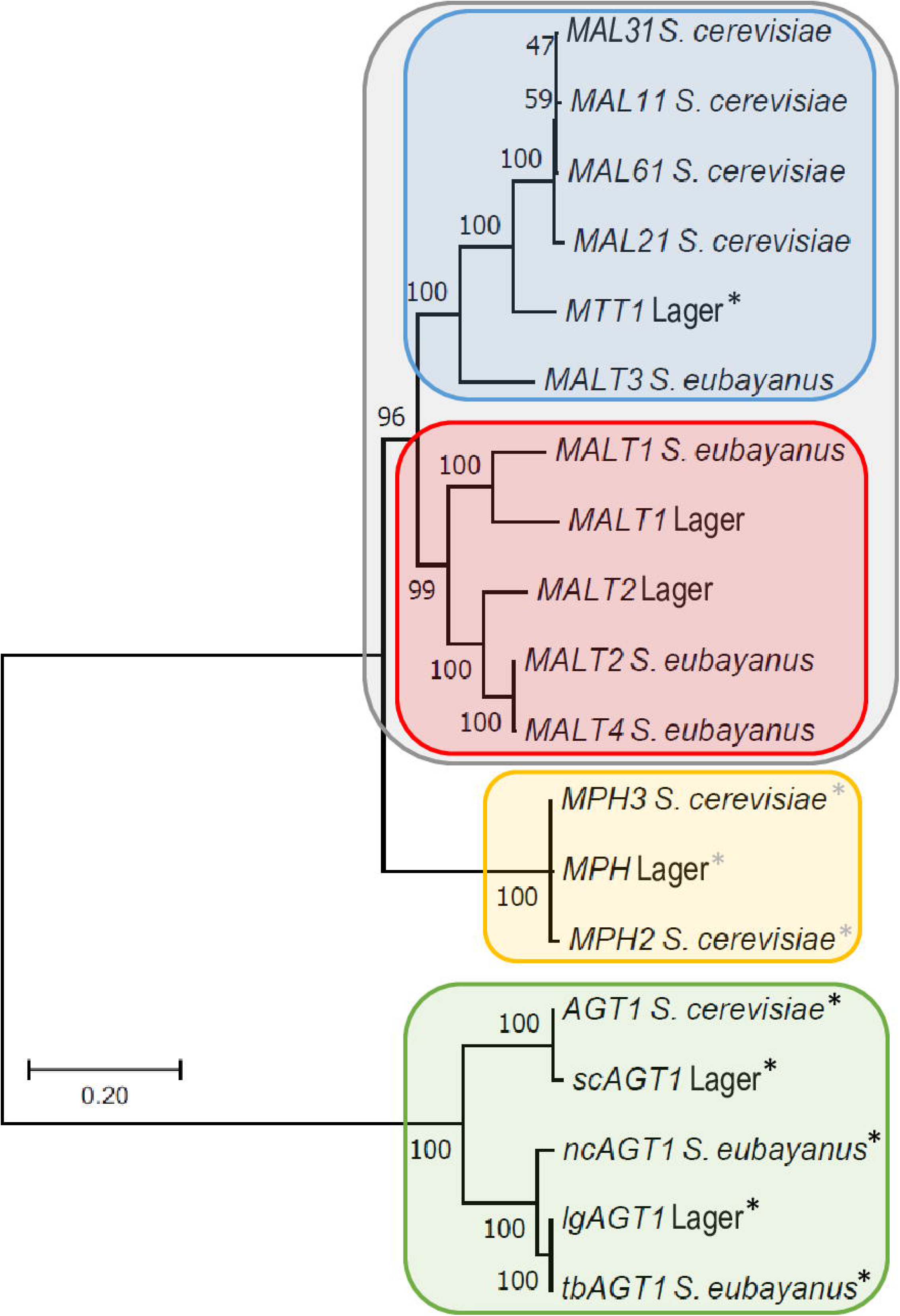
Phylogeny of *Saccharomyces MALT* genes. ML phylogenetic tree of *MALT* genes described in *S. cerevisiae, S. eubayanus*, and lager-brewing hybrids. The scale bar equals the number of nucleotide substitutions per site. Black “*” indicate genes characterized as encoding proteins capable of transporting maltotriose. Gray “*” indicates genes encoding transporters whose ability to transport maltotriose is ambiguous.

### Indirect evolution of maltotriose consumption

Since yHRVM108 contains a functional maltotriose transporter whose overexpression is sufficient for growth on maltotriose (Table 1), we anticipated that it would be simple for yHRVM108 to evolve the ability to utilize maltotriose under direct selection for this trait. Because yHRVM108 is unable to grow on maltotriose, we passaged the strain in 2% maltotriose medium with a small amount of added glucose to permit a limited number of cell divisions to allow for mutation and selection to occur. Over the course of 100 passages under this selection regime, representing around 1,050 cell divisions across three experimental replicates, no maltotriose-utilizing lineage of yHRVM108 arose.

While evolving yHRVM108 directly for maltotriose consumption was not successful, we were initially surprised to find an alternative and indirect selection regime was effective at evolving maltotriose utilization in this background. When we began adaptive evolution of yHRVM108 on maltotriose, we also began selecting for increased growth of yHRVM108 on maltose to try and improve this strain’s sluggish growth on this carbon source. All three replicates of this experiment eventually evolved the ability to grow rapidly on maltose. Interestingly, in addition to growing on maltose four times more rapidly over two days (S2 Table), single-colony isolates from the first two replicates that evolved rapid maltose utilization also gained the ability to utilize maltotriose (S3 Table), despite never being exposed to maltotriose during the course of the adaptive evolution experiment. The fact that maltotriose consumption independently evolved during selection for improved maltose utilization multiple times, most likely through increased expression of *ncAGT1*, suggests that our maltotriose selection regime itself may have played a role in restraining evolution.

Though we found the difficulty of evolving expression of a functional transporter surprising, such a result is not unprecedented. In a long-term evolution experiment in *Escherichia coli*, a functioning citrate transporter was present in the founding strain. Though expression of this gene would have been highly favored in the citrate-rich experimental environment, it took thousands of generations, even after the necessary potentiating mutations had appeared, before a gene amplification and rearrangement event joined the citrate transporter gene to a new promoter, resulting in a novel expression pattern (70). These results show how an organism’s preexisting genetic architecture, interacting with the selective environment, can facilitate or impede evolution along a particular path (71–74). Since maltose is consumed to some extent by yHRVM108, loosened regulation of *ncAGT1* might have been selected for alongside other *MAL* genes because *AGT1-*type transporters have a broad substrate range that includes maltose (30). In contrast, evolving maltotriose utilization directly might have required specific and rare changes to the *ncAGT1* locus itself. In retrospect, what appeared to be a simple request, to turn on the *ncAGT1* gene in the presence of maltotriose as the sole carbon source, may in fact have been quite difficult by simple mutations, whereas our indirect selection regime on maltose proved more effective.

### Evolution of maltotriose utilization through a chimeric transporter

To determine how strains lacking any maltotriose transporters could evolve them, we also tried to experimentally evolve maltotriose utilization in the *S. eubayanus* strains FM1318 (48) and in yHKS210 (53). A search of the available genome sequence reads for FM1318 (54,59) and yHKS210 (52) confirmed that neither of these strains contain genes that are closely related to *AGT1-*like genes or other known maltotriose transporters (28,29). Based on our analysis of the available whole-genome sequencing data, these strains only contain the four *MALT* transporter genes previously identified in FM1318 (54), which are unable to confer maltotriose utilization even when overexpressed (Table 1). Since, like yHRVM108, neither of these strains could grow on maltotriose, a small amount of glucose was also added to the medium to permit a limited number of cell divisions for mutation and selection. Over the course of 100 passages, representing approximately 2,100 cell divisions in total between the two strains and their replicates, a single replicate, derived from strain yHKS210, evolved the ability to grow on maltotriose. Two single-colony isolates (yHEB1505-6) from this replicate were isolated and confirmed to be able to grow on maltotriose without added glucose (Fig 3A, S3 Table).

**Fig 3.**
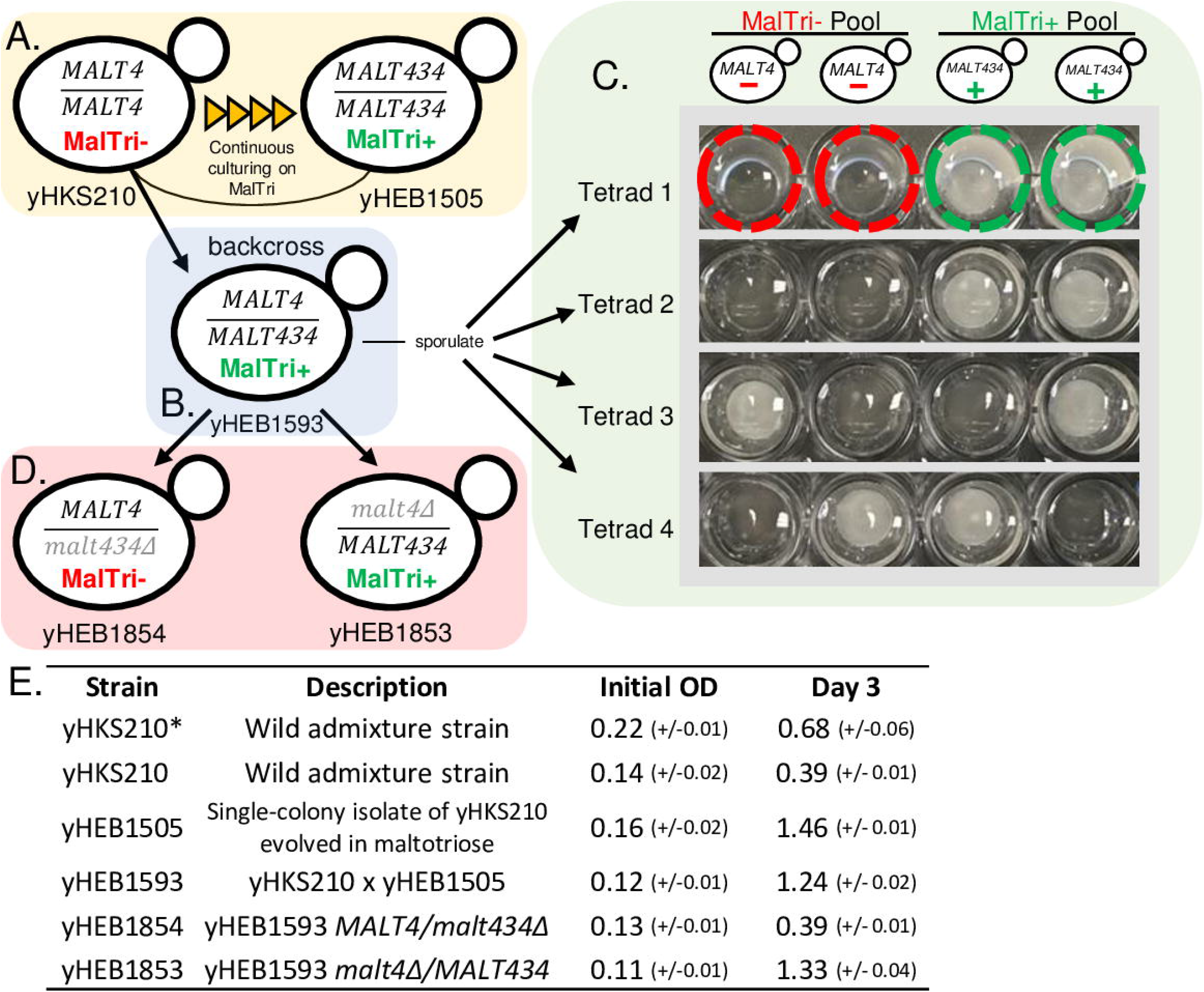
Evolution and validation of the chimeric maltotriose transporter Malt434. A) After continuous culturing on maltotriose with a small amount of added glucose, yHKS210, which was originally unable to use maltotriose (MalTri−), evolved the ability to consume maltotriose (MalTri+). B) Strain yHEB1593, which is a backcross between yHKS210 and yHEB1505, was also MalTri+. C) To test the inheritance of maltotriose utilization, yHEB1593 was sporulated. The panel shows a subset of tetrads screened growing on SC + 2% maltotriose. Examples of MalTri-spores in Tetrad 1 are circled in red, and MalTri+ examples are circled in green. Whole genome sequencing of MalTri+ and MalTri-pools showed that maltotriose utilization perfectly correlated with the presence/absence of *MALT434*. D) Reciprocal hemizygosity test (78) of the *MALT4/MALT434* locus in the backcross strain yHEB1593. E) Table of initial and day-three OD_600_(OD) readings of yHKS210, yHEB1505, yHEB1593, yHEB1853, and yHEB1854 on SC + 2% maltotriose as the sole carbon source. N = 3, standard deviation in parentheses. * Control grown in SC + 0.04% glucose to reflect the approximate amount of growth expected from contamination with other carbon sources when using 98% pure maltotriose.

To determine the genetic architecture of maltotriose utilization in the replicate of yHKS210 that evolved the ability to grow on maltotriose, we set up an F_1_ backcross between the evolved maltotriose-utilizing isolate yHEB1505 and the parent strain (yHKS210), producing strain yHEB1593, a putative heterozygote capable of growth on maltotriose (Fig 3B and E). In a test of 15 fully viable F_2_ tetrads, maltotriose utilization segregated in a perfect 2:2 manner (Fig 3C). These results suggest that the ability of the evolved strain to utilize maltotriose is conferred by a dominant mutation at a single genetic locus. We performed bulk segregant analysis (75–77) using strains derived from the F_2_ spores, dividing them between those that could (MalTri+) and those that could not (MalTri−) utilize maltotriose (Fig 3C), with a total of 30 strains in each category. Twelve 1-kb regions were identified as potentially containing fixed differences between the MalTri+ and MalTri− strains. Of these regions, eight mapped to genes encoding ribosomal proteins and most likely represent assembly artefacts due to the presence of many closely related paralogs and/or their absence from the MalTri− de novo assembly that was used for comparisons. Three other regions contained fixed differences between the MalTri+ and MalTri− groups but had no clear relationship to carbon metabolism. The final 1-kb region mapped to the *MALT4* locus of *S. eubayanus* genome (54,59). The coding sequence of *MALT4* from the MalTri+ group contained 52 SNPs relative to the *MALT4* allele found in yHKS210, all of which occurred within a single 230-bp region. Of these, 11 were predicted to lead to non-synonymous changes. Closer inspection revealed that the changes within the 230-bp region were the result of a recombination event between *MALT4* and *MALT3*, creating a chimeric gene (Fig 4), likely through ectopic gene conversion. We call this chimeric *MALT4* allele *MALT434* after the arrangement of sequences from its parent genes. The sequence of *MALT3* was not impacted by this mutational event.

**Fig 4.**
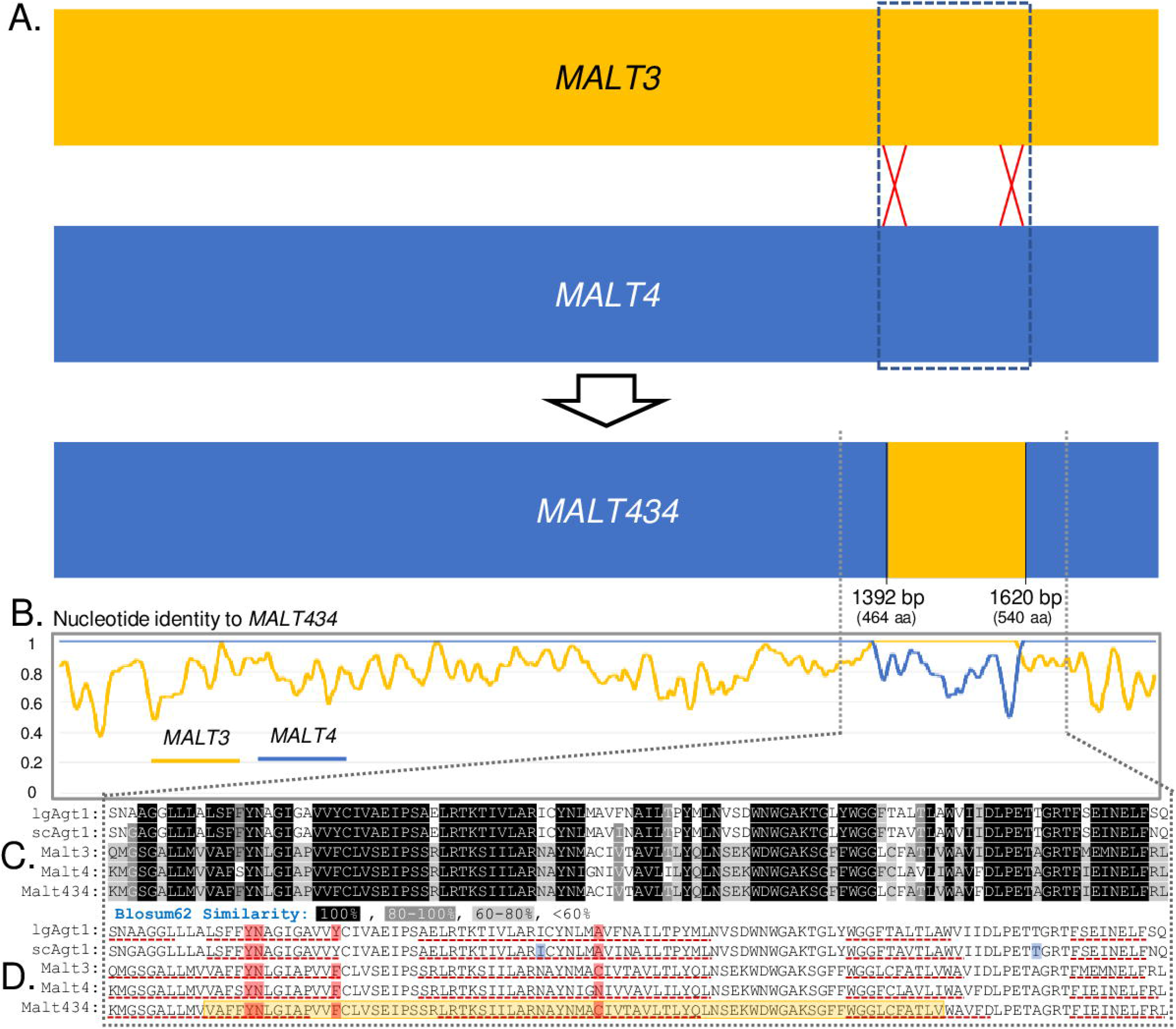
Sequence architecture of *MALT434*. A) Schematic of the origin of *MALT434*. B) Line graphs representing the identity between nucleotide sequences of *MALT3* and *MALT4* from yHKS210 to *MALT434* over 10-bp sliding windows. C-D) Segment of the alignment of the chimeric region between Malt3, Malt4, Malt434, scAgt1, and Igagt1. The region highlighted in yellow in the Malt434 sequence indicates the chimeric region. The regions underlined with a red dashed line are predicted transmembrane domains. The amino acids highlighted in red are predicted maltose-binding residues. The residues highlighted in blue were experimentally found to be important for maltotriose transport by Smit *et al*. 2008.

To confirm that *MALT434* was the causative locus of maltotriose utilization, we performed a reciprocal hemizygosity test (78) in the heterozygous F_1_ backcross strain (Fig 3D). Removal of *MALT434* eliminated the F_1_ backcross strain’s ability to utilize maltotriose (Fig 3E), demonstrating that *MALT434* is required for maltotriose utilization. Conversely, removing the parental, non-chimeric allele of *MALT4* in the heterozygous F_1_ backcross strain had no impact on maltotriose utilization. Furthermore, overexpression of Malt434 in both the unevolved parent, yHKS210, and in the yHRVM108 background (Fig 5) supported growth on maltotriose, demonstrating that overexpression of Malt434 is sufficient to confer maltotriose utilization. These results strongly suggest that the mutant *MALT434* gene encodes a functional maltotriose transporter.

**Fig 5.**
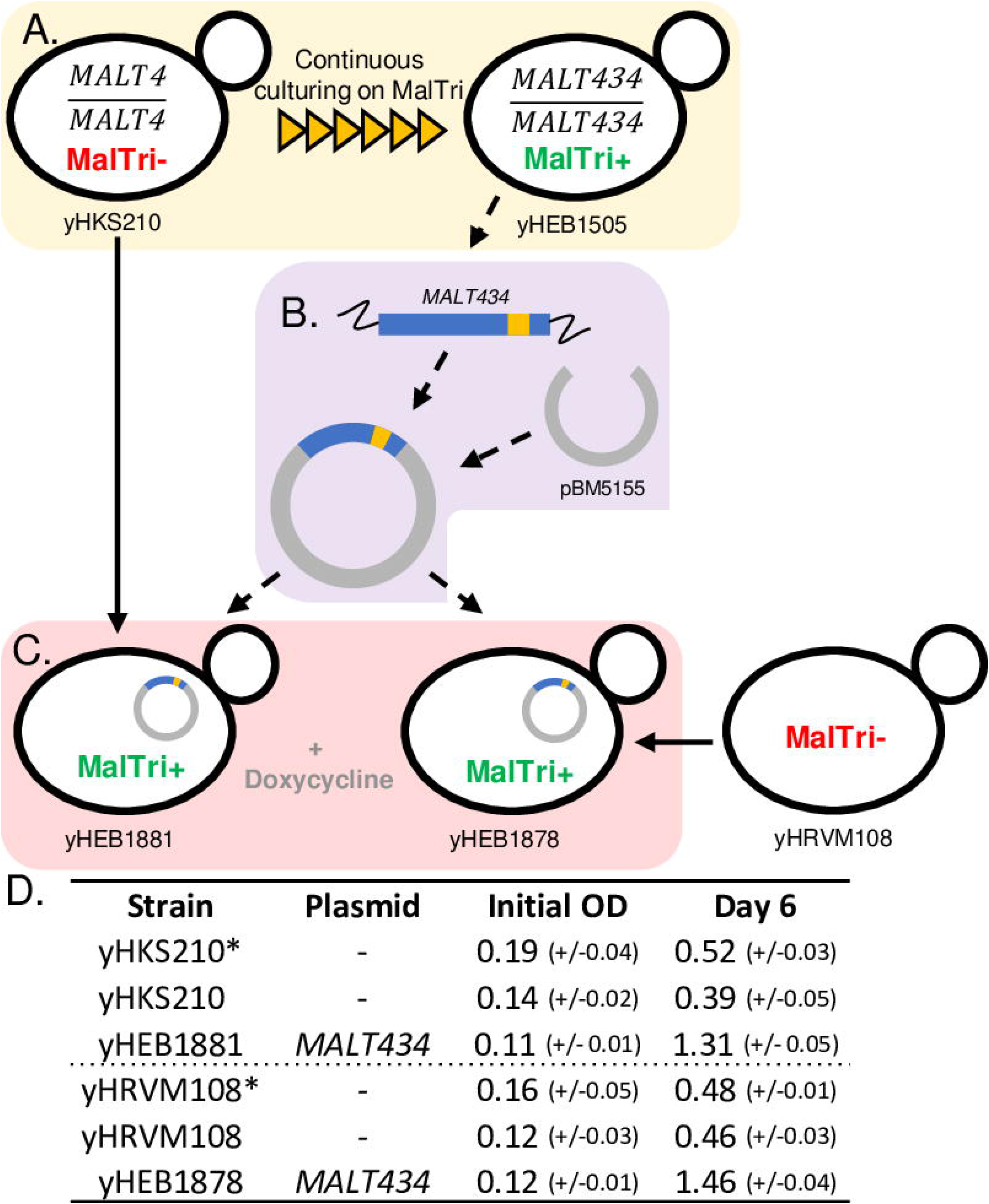
Heterologous expression of *MALT434*. A) Evolution of non-maltotriose utilizing strain (MalTri−), yHKS210, to maltotriose utilizing (MalTri+) strain, yHEB1505, by serial passing on maltotriose containing media (same as Fig 3A). B) Insertion of *MALT434* into vector pBM5155 for doxycycline-inducible heterologous expression in MalTri− strains. C) Transformation of *MALT434* expression plasmid in MalTri− *S. eubayanus* strains yHKS210 and yHRVM108. D) Table of initial and day-six OD_600_ (OD) measurements of parent strains and strains carrying the *MALT434* expression plasmid grown in SC media with maltotriose as the sole carbon and doxycycline to induce plasmid expression. N = 3, standard deviation in parentheses. * Control grown in SC + 0.04% glucose + doxycycline to reflect the approximate amount of growth expected from contamination with other carbon sources when using 98% pure maltotriose.

### Potential structural impact of Malt434 chimerism

It was surprising that sequences from *MALT3* enabled *MALT4* to encode a maltotriose transporter because neither *MALT3* nor *MALT4* supported maltotriose utilization on their own (Table 1). Malt3 and Malt4 share about 80% amino acid identity overall and 85% amino acid identity in the chimeric region specifically (Fig 4B). Most residues in the chimeric region had high similarity between Malt3 and Malt4, as measured by Blosum62 similarity matrix (Fig 4C) (79), but there were a handful of low-similarity amino acids as well. To gain insight into what changes in protein structure may be driving the new functionality of Malt434, we used I-TASSER (80–82) to predict the protein structure of Malt3, Malt4, and Malt434. I-TASSER predicts a protein’s structure based on its homology to proteins whose structures have already been solved. Consistent with other studies on the structure of maltose transporters in *Saccharomyces* (27,83–85), I-TASSER predicted that Malt3, Malt4, and Malt434 were similar to members of the Major Facilitator Superfamily (MFS) of transporters, specifically the sugar porter family (85). Protein structure is predicted to be conserved between Malt3 and Malt4, including within the chimeric region, which encompasses one full transmembrane domain and parts of two other transmembrane domains (Fig 4D). Four maltose-binding sites were also predicted in the chimeric region. These same domains and predicted binding residues were predicted for Malt434 as well. Interestingly, I-TASSER predicted several of the alpha helices to be shorter in the chimera relative to the parent proteins: two alpha helices in the chimeric region and two towards the N-terminal end of the protein (Fig 4D, S1 Fig). The regions covered by these alpha helices were otherwise predicted to be conserved, out to phylogenetically distantly related Malt proteins IgAgt1 and scAgt1 (Fig 2, Fig 4D, S1 Fig). The predicted shortening of some alpha helices suggests that recombining the *MALT3* region into *MALT4* may have decreased the overall rigidity of the encoded chimeric protein, allowing it to accommodate bulkier substrates, such as maltotriose. Mutations that increase structural flexibility have been recognized in protein engineering as an important step in accommodating new substrates (86,87).

Besides increasing overall flexibility, the specific location of the chimeric region could have also played a role in supporting maltotriose transport. A previous study found that two residues were important for scAgt1’s ability to transport maltotriose, while not affecting its ability to transport maltose (44). One of these residues lies within the chimeric region of Malt434, and the other is 10 amino acids downstream (Fig 4D, S1 Fig). Since the overall structure of maltose/maltotriose transporters is conserved (27,83–85), the area in and around the chimeric region of Malt434 may itself be important for substrate specificity.

Thus, the chimeric structure of Malt434 may have facilitated maltotriose transport in two ways. First, it may have increased the overall flexibility of the protein, allowing it to accommodate the larger maltotriose molecule. Second, it could also have specifically altered an important substrate interface to facilitate a better interaction with maltotriose, possibly also by making this region more flexible. Testing these biophysical and structural models will require future experiments, such as solving the crystal structures for Malt3, Malt4, and Malt434 as complexes with maltose and/or maltotriose.

### A non-modular chimeric path to novel substrate utilization

Most of the work on functional innovations by chimeric proteins has focused on the rearrangement of discrete functional units, with or without the benefit of intronic sequences (6,7,14,88–91). However, Malt434 does not fit easily into the framework of new protein creation by the reordering/exchanging of modules, even when considering smaller functional units such as a single alpha helix. While the chimeric region does completely move one alpha helix from Malt3 into the Malt4 background, the breakpoints of the gene conversion also result in two other alpha helices with some residues from the Malt4 parent and some from the Malt3 parent, creating chimeric alpha helices (Fig 4D, S1 Fig). In addition, while domains important for sugar specificity probably exist in Malt3 and Malt4 (44,84), with respect to maltotriose, the “sugar specificity” domain(s) between Malt3 and Malt4 do not seem to have different functions or specificities in their native backgrounds. In Malt3 and Malt4, there are no specific “maltotriose-transporting” domains to be swapped. Instead, the ability of the residues from Malt3 to facilitate maltotriose transport likely relies on their interaction with one or more residues in Malt4, not on their independent ability to interact with maltotriose.

Rather than the modular framework of novel protein formation, Malt434 exemplifies another framework for how recombination can lead to the evolution of novel functions. Theoretical and experimental work has demonstrated the important role that recombination between related proteins can play in facilitating the evolution of new functions (3,4,15,92). Indeed, protein engineering has utilized the technique of DNA shuffling since the mid-1990’s to recombine closely related coding sequences to efficiently generate proteins with novel or improved functions (19). More recently, experimental work has begun to demonstrate the importance of recombination between closely related proteins in nature for the evolution of new functions (15,18,92). In this model, two duplicate proteins neutrally accumulate the multiple amino acid changes needed for a new function independently. All of the mutations that are needed for the new function are then brought together at once, en masse, through recombination. This molecular mechanism allows proteins to “tunnel” to new functions, bypassing potentially deleterious intermediates that would be encountered through a series of amino acid substitutions (3,4).

While *MALT3* and *MALT4* are not recent duplicates, they are distant paralogs (Fig 2). In addition, as members of the sugar porter subfamily of proteins, they share a highly conserved protein structure (27,83–85). The conservative nature of sugar porter family proteins means that recombination events like the one that formed Malt434, which do not fall between clear domains, probably have a relatively high likelihood of creating functional transporters (93), albeit ones of unpredictable specificity. In the case of Malt434, we do not yet know which specific amino acid interactions were important for the gain of maltotriose utilization in the chimera, let alone the function or history of the residues in the background of their native protein sequences. It may be that they represent neutral changes in their parental backgrounds, but they also could have been selected for other specificities. Nevertheless, the independent accumulation of these changes in a common ancestral protein background eventually allowed these sequences to recombine and create a novel function.

## Conclusions

Our findings suggest that the evolution of maltotriose utilization by *Saccharomyces* yeasts is not a straightforward process. Even when a functioning maltotriose transporter is available in the parent genome, the regulatory changes necessary to support atypical expression may be difficult to evolve under certain experimental conditions. Conversely, when a maltotriose transporter is not already present, single point mutations are probably insufficient to switch or expand the specificity of available Malt proteins. Recombination between paralogous proteins can rapidly do what a single point mutation cannot and, in a single rare mutational event, introduce the multiple residue changes needed to perform a new function. Our report on the evolution of a chimeric maltotriose transporter from parental proteins that could not transport maltotriose supports the role of recombination, beyond the simple swapping of functional protein domains and peptide motifs, in the formation of proteins with novel functions.

## Materials and Methods

### Strains

All strains discussed in this paper are listed in S4 Table. Briefly, FM1318 is a monosporic derivative of the type strain of *S. eubayanus*, which was isolated from Patagonia (48). yHRVM108 was isolated from Durham, North Carolina, and is closely related to the *S. eubayanus* strains that hybridized with *S. cerevisiae* to give rise to lager-brewing yeasts (52). yHKS210 was isolated from Sheboygan, Wisconsin, and is the result of admixture between populations A and B of *S. eubayanus*. yHKS210 is nearly homozygous due to selfing after the initial admixture event (53). Of these strains, FM1318 and yHKS210 grew well on maltose, but they did not grow on maltotriose. yHRVM108 grew sluggishly on maltose and did not grow on maltotriose. yHAB47 is a copy of Weihenstephan 34/70 (52), a representative of the Frohberg or Group II (94) lineage of lager-brewing hybrids (S. *cerevisiae* (2n) × *S. eubayanus* (2n) (58)). CDFM21L.1 is a strain of *S. eubayanus* isolated from Tibet (51) and is closely related to yHRVM108. Of known *S. eubayanus* strains, CDFM21L.1 is the most genetically similar to the *S. eubayanus* parents of lager-brewing hybrids (51,52).

### Identification of *MALT* genes

Previously, we identified four genes with homology to genes encoding maltose transporters in *S. cerevisiae* and lager-brewing hybrids in the genome assembly of FM1318 published by Baker *et al*. 2015 (54). These genes were previously designated *MALT1-4*. Only a partial contig was available for *MALT4* in this assembly, but a BLAST (95) search of the Okuno *et al*. 2016 (59) assembly of the type strain of *S. eubayanus* (of which FM1318 is a monosporic derivative) allowed us to annotate the full-length sequence of *MALT4. MALT4* has 99.7% identity to *MALT2* at the nucleotide level and is predicted to have 100% identity at the amino acid level. The regions from 900 bp downstream of *MALT2* and *MALT4* and upstream to the ends of chromosomes V and XVI (regions of approximately 12 kb in the Okuno *et al*. 2016 (59) assembly), respectively, share 99.1% nucleotide identity. The 10 kb outside of this region only share 49.8% nucleotide identity. Thus, *MALT2* and *MALT4* are close paralogs that are likely related by a recent subtelomeric duplication and/or translocation event.

Reads for homologs of *AGT1* were retrieved using the functional *AGT1* sequence from lager yeast (*lgAGT1*) as the query sequence (55) in an SRA-BLAST search of the SRA databases of NCBI for yHRVM108 (SRR2586159) and CDFM21L.1 (SRR1507225). All reads identified in the BLAST searches were downloaded and assembled using the de novo assembler in Geneious v. 9.0.3 (http://www.geneious.com) (96). The homologs identified in yHRVM108 and CDFM21L.1 were designated *ncAGT1* (for North Carolinian *AGT1*) and *tbAGT1* (for Tibetan *AGT1*), respectively. The presence and sequence of *ncAGT1* in yHRVM108 was further verified by PCR amplification and Sanger sequencing (S5 Table). CDFM21L.1 was not available at the time of this work for further verification of the presence of *tbAGT1*.

### Adaptive evolution

Design of the adaptive evolution experiments was based on Parreiras *et al*. 2014 (97). The highest available purities of carbon sources were used: 98% pure maltotriose, ≥99% pure maltose, and 100.0% pure glucose. Adaptive evolution was initiated by growing parent strains overnight in liquid YPD medium (1% yeast extract, 2% peptone, 2% glucose). One mL of maltotriose or maltose medium was inoculated with enough overnight culture to give an OD_600_ reading of ~0.1, as measured with an IMPLEN OD600 DiluPhotometer™. Evolution on maltotriose was conducted in synthetic complete (SC) medium (0.17% yeast nitrogen base, 0.5% ammonium sulfate, 0.2% complete drop out mix) with 2% maltotriose and 0.1% glucose. The addition of 0.1% glucose ensured enough growth that mutations could occur and be selected for through the ensuing generations. Adaptive evolution of yHRVM108 on maltose was carried out in SC with 2% maltose. Because yHRVM108 grew so poorly on maltose alone, an additional 0.1% glucose was supplemented into its medium; after increased growth was observed around generation 110 for replicate A (from which strains yHEB1585-1587 were derived), around generation 80 for replicate B (from which strains yHEB1588-90 were derived), and around generation 155 for replicate C (from which strains yHEB1778-80 were derived), subsequent generations of yHRVM108 adaptive evolution on maltose for these replicates were conducted with 2% maltose only. Adaptive evolution experiments of each strain were carried out in triplicate. Samples were grown on a culture wheel at room temperature (22°C) and diluted 1:10 into fresh media every 3-4 days. Samples of each evolution replicate were taken every other passage and placed into long-term storage by mixing 700uL of culture with 300uL of 50% glycerol in a cryotube and storing it at −80°C. The numbers of doublings between passages were estimated from cell counts during the second and third passages. Evolution was carried out for a total of 100 passages. Strains that could not use the primary carbon source in the adaptive evolution medium underwent approximately one cell division per day on average.

### Sporulation and backcrossing

To induce sporulation, strains were grown to saturation, washed twice, and then resuspended in 200μL liquid sporulation (spo) medium (1% potassium acetate, 0.5% zinc acetate). 30μL of this suspension was added to 1.5mL of spo medium and incubated on a culture wheel at room temperature. Cultures were checked for sporulation after 2–5 days. Tetrads were dissected using a Singer SporePlay. For backcrossing, tetrads of the strains to be crossed were dissected on a single YPD plate. A spore from one parent was placed in close proximity to a spore from the other parent, and they were observed over several hours for mating and zygote formation. Transformations of the diploid F_1_ backcross strain for gene knockouts were carried out as described below in the section describing the construction of gene expression plasmids.

### Construction of gene expression plasmids

Genes encoding transporters of interest were cloned via gap repair into the *Not*I site of plasmid pBM5155 (GenBank KT725394.1), which contains the complete machinery necessary for doxycycline-based induction of genes cloned into this site (98). Transformation was carried out using standard lithium acetate transformation (99) with modifications to optimize transformation in *S. eubayanus*. Specifically, transformation reactions were heat-shocked at 34°C. After 55 minutes, 100% ethanol was added to 10% total volume, and the reactions heat shocked for another 5 minutes before they were allowed to recover overnight and plated to selective media the next day. When necessary, plasmids were recovered and amplified in *E. coli* for transformation into multiple strains. The sequences of genes encoding transporters cloned into pBM5155 were verified by Sanger sequencing. *S. eubayanus MALT1, MALT3*, and *MALT4* were amplified from FM1318, *IgAGT* was amplified from yHAB47, and *ncAGT1* was amplified from yHRVM108. Primers used for plasmid construction and sequence verification are listed in S5 Table.

### Growth assays

Growth was measured in liquid media in 96-well plates using OD_600_ measurements on a FLUOstar Omega^®^ microplate reader. Strains were first grown to saturation in liquid YPD medium, then washed twice and diluted in SC without added carbon to OD_600_ = 1.9 +/− 0.05 to ensure that all cultures had approximately the same starting concentration. 15μL of each diluted culture was added to 235μL of the test medium. Three technical replicates, randomly distributed on a 96-well plate to control for position effects, were carried out for each strain. Single-colony isolates of yHKS210 evolved on maltotriose and single-colony isolates of yHRVM108 evolved on maltose were tested in SC medium + 2% maltotriose. Single-colony isolates of yHRVM108 evolved on maltose were also tested on SC medium + 2% maltose. Strains carrying *MALT* genes expressed on an inducible plasmid were tested in SC medium + 2% maltotriose and 5 ng/mL doxycycline to induce plasmid gene expression. To control for growth from the small amount of non-maltotriose sugar in 98% pure maltotriose, the parent strains of yHRVM108 and yHKS210 were also tested in SC medium + 0.04% glucose, reflecting the approximate amount of other carbon sources expected in SC medium + 2% maltotriose.

### Bulk segregant analysis

60 spores from 15 fully viable tetrads of strain yHEB1593 (F_1_ of yHKS210 × yHEB1505) were dissected and individually screened for their ability to grow in SC + 2% maltotriose. F_2_ segregants that could grow on maltotriose were classified as MalTri+, and those that could not were classified as MalTri−. Each F_2_ segregant was then individually grown to saturation in liquid YPD. The saturated cultures were spun down, the supernatant removed, and enough cells resuspended in liquid SC medium to give an OD_600_ measurement of between 1.9 and 1.95, as measured with an IMPLEN OD600 DiluPhotometer™. Strains were pooled based on their ability to grow on maltotriose, forming a MalTri+ pool and a MalTri− pool. To pool, 1mL of each strain dilution was added to the appropriate pool of cells and vortexed to mix. Phenol-chloroform extraction and ethanol precipitation was used to isolate gDNA from the segregant pools. The gDNA was sonicated and ligated to Illumina TruSeq-style dual adapters and index sequencing primers using the NEBNext^®^ DNA Library Prep Master Mix Set for Illumina^®^ kit following the manufacturer’s instructions. The paired-end libraries were sequenced on an Illumina MiSeq instrument, conducting a 2 × 250bp run.

### Analysis of bulk segregant sequencing reads

To identify fixed differences between the meiotic segregant pools, de novo assemblies were made for the MalTri− group of segregants using the meta-assembler iWGS with default settings (100). The final genome assembly of the MalTri− pool was made by DISCOVAR (101) in iWGS. This assembly was used for reference-based genome assembly and variant calling using reads from the MalTri+ pool following the protocol described in Peris and Langdon *et al*. 2016 (52). Assemblies of the putative chimeric maltotriose transporter were retrieved from the MalTri+ pool of reads using the program HybPiper (102). Briefly, HybPiper uses a BLAST search of read sequences to find reads that map to a query sequence; it then uses the programs Exonerate (103) and SPAdes (104) to assemble the reads into contigs. The sequence and genomic location of the chimeric transporter were further verified by PCR amplification and Sanger sequencing (S5 Table), as was the sequence of *MALT4* from yHKS210.

### Phylogenetic analyses and computational predictions of protein structures and functions

Multiple sequence alignments between the proteins encoded by the *MALT* genes were carried out using MUSCLE (105), as implemented in Geneious v.9.0.3 (96) (http://www.geneious.com). Phylogenetic relationships were determined using codon alignments. Codon alignments were made using PAL2NAL (Suyama, Torrents, & Bork, 2006; http://www.bork.embl.de/pal2nal/) to convert the MUSCLE alignments of amino acid sequences to nucleotide alignments. A phylogenetic tree of nineteen *MALT* genes from *S. eubayanus* and *S. cerevisiae* and lager-brewing yeasts was constructed as described in Baker *et al*. 2015 (54) using MEGA v.6. Most genes used in the phylogenetic analysis were retrieved as previously described in Baker *et al*. 2015 (54) as follows: *MAL21, MAL31*, and *MAL61* from *S. cerevisiae; MALT1* and *MALT3* from *S. eubayanus; MALT1, MALT2*, and *MPH* from lager-brewing yeast; *MPH2 and MPH3* from *S. cerevisiae; AGT1* (*MAL11* in Baker *et al*. 2015 (54)) from *S. cerevisiae; scAGT1* (*WeihenMAL11-*CB in Baker *et al*. 2015 (54)); and *lgAGT1* (*WeihenMAL11-*CA in Baker *et al*. 2015 (54)) Sequences for *MALT2* and *MALT4* were retrieved from the genome assembly of CBS 12357^T^ from Okuno *et al*. 2016 (59). *MAL11* was retrieved from the genome assembly of *S. cerevisiae* strain YJM456 (107). Sequences for *tbAGT1* and *ncAGT1* were retrieved as described above. *MAL11* and *AGT1* both encode a-glucoside transporters located at the *MAL1* locus in *S. cerevisiae* and, as such, are considered alleles of each other (27,108). Their shared genomic location notwithstanding, *MAL11* and *AGT1* are not phylogenetically closely related, with *MAL11* clustering with other *MALx1* type transporters (Fig 2). In addition, while *AGT1* can support maltotriose transport, *MAL11*, like other known *MALx1* genes, cannot (27,30). Despite their dissimilarity, *AGT1* is recorded in the *Saccharomyces* Genome Database (yeastgenome.org) as *MAL11* since the reference strain carries the *AGT1* allele at the *MAL1* locus (60,63). For this reason, *MAL11* is often used to refer to *AGT1* (30,32,54). For clarity, here we use *MAL11* to only refer to the *MALx1-*like allele and *AGT1* to refer to the distinct maltotriose-transporting allele.

Protein structure predictions for *MALT3, MALT4, IgAGT1*, and *scAGT1* were carried out using the I-TASSER server, and the structure prediction of *MALT434* was carried out using the command line version of I-TASSER (80–82) (https://zhanglab.ccmb.med.umich.edu/l-TASSER/, accessed between 2-7-2018 and 2-28-2018). The potential impact of the single residue difference between *IgAGT1* and *tbAGT1* was analyzed by two different methods. Prediction of the change in free energy (ΔΔ*G*) was carried out using the STRUM server (https://zhanglab.ccmb.med.umich.edu/STRUM/, accessed 3-21-18) (65). A ΔΔG score of < +/− 0.5 was considered to be unlikely to affect function (109). Homology-based predictions were made using SIFT at http://sift.icvi.org/ (accessed 3-30-18) (66,110–113). The SIFT Related Sequences analysis was done using the amino acid sequences of *MALT* genes in the phylogenetic analysis above. Several SIFT analyses were also carried out using the SIFT Sequence analysis program. This analysis operates using the same principle as the SIFT Related Sequences analysis, but rather than being supplied by the user, homologous sequences were provided by a PSI-BLAST search of the indicated protein database. The SIFT Sequence analyses were carried out using default settings and the following databases available on http://sift.icvi.org/ (accessed 3-30-18): NCBI nonredundant 2011 Mar, UniRef90 2011 Apr, UniProt-SwissProt 57.15 2011 Apr.

## Acknowledgments

We thank Dana A. Opulente, Quinn K. Langdon, Ryan V. Moriarty, and Kelly V. Buh for their assistance maintaining the experimental evolution lines described in this manuscript.

## Supporting Information

**S1 Fig. Protein structural alignment between Malt3, Malt4, Malt434, scAgt1, and Igagt1.** Protein structural alignment between Malt3, Malt4, Malt434, scAgt*1*, and IgAgt*1*. The purple blocks represent predicted alpha helices, and the orange lines represent predicted beta strands. Red ticks mark predicted maltose-binding sites. Blue ticks mark residues found to be important for maltotriose transport by Smit *et al*. 2008. A green tick marks the location of the single non-synonymous substitution between *IgAGT1* and *tbAGT1*. Arrows point to alpha helices in Malt434 whose predicted sizes are reduced compared to other transporters in the alignment.

**S1 Table. Mutation prediction analyses.** Results of mutation prediction analyses for E18V, the sole amino acid substitution in the IgAgt*1* protein sequence, relative to tbAgt1.

**S2 Table. Maltose growth assay.** Growth on maltose of single-colony isolates. Isolated from adaptive evolution of yHRVM108 on 2% maltose + 0.1% glucose. N = 3. * Control grown in SC + 0.04% glucose to reflect the approximate amount of growth expected from contamination with other carbon sources when using 98% pure maltotriose.

**S3 Table. Maltotriose growth assay.** Growth on maltotriose of single-colony isolates from adaptive evolution experiments. Strains were evolved with either maltotriose or maltose as the primary carbon source (2%) with 0.1% added glucose. N = 3. * Control grown in SC + 0.04% glucose to reflect the approximate amount of growth expected from contamination with other carbon sources when using 98% pure maltotriose.

**S4 Table. Strains and plasmids used in this work.**

**S5 Table: Oligonucleotides used in this work.**

